# Modeling *IKZF1* lesions in B-ALL reveals distinct chemosensitivity patterns and potential therapeutic vulnerabilities

**DOI:** 10.1101/2020.05.26.109579

**Authors:** Jason H. Rogers, Rohit Gupta, Jaime M. Reyes, Michael C. Gundry, Geraldo Medrano, Anna Guzman, Tidie Song, Cade Johnson, Sean Barnes, Carlo D. D. Cristobal, Lorenzo Brunetti, Margaret A. Goodell, Rachel E. Rau

## Abstract

IKAROS family zinc finger 1 (*IKZF1*) alterations represent a diverse group of genetic lesions that are associated with an increased risk of relapse in B-lymphoblastic leukemia (B-ALL). Due to the heterogeneity of concomitant lesions it remains unclear how *IKZF1* abnormalities directly affect cell function and therapy resistance and whether their consideration as a prognostic indicator is valuable in improving outcome. We used CRISPR/Cas9 to engineer multiple panels of isogeneic lymphoid leukemia cell lines with a spectrum of *IKZF1* lesions in order to measure changes in chemosensitivity, gene expression, cell cycle, and in vivo engraftment dynamics that can be directly linked to loss of IKAROS protein. *IKZF1* knockout and heterozygous null cells displayed relative resistance to a number of commonly employed therapies for B-ALL including dexamethasone, vincristine, asparaginase, and daunorubicin. Transcription profiling revealed a stem/myeloid cell-like phenotype and JAK/STAT upregulation after IKAROS loss. We also used a CRISPR homology-directed repair (HDR) strategy to knock-in the dominant-negative IK6 isoform tagged with GFP into the endogenous locus and observed a similar drug resistance profile with the exception of retained sensitivity to dexamethasone. Interestingly, *IKZF1* knockout and IK6 knock-in cells both have significantly increased sensitivity to cytarabine, suggesting intensification of nucleoside analog therapy may be specifically effective for *IKZF1*-deleted B-ALL. Both types of *IKZF1* lesions decreased survival time of xenograft mice, with higher numbers of circulating blasts and increased organ infiltration. Given these findings, exact specification of *IKZF1* status in patients may be a beneficial addition to risk stratification and could inform therapy.

**Key points:**

- Engineered *IKZF1* perturbations result in a stem-cell like expression signature, enhanced engraftment in vivo, and multi-drug resistance
- Loss of IKAROS may result in new vulnerabilities due to increased sensitivity to cytarabine and upregulation of JAK/STAT and mAb targets

## Introduction

Approximately 20% of pediatric patients and the majority of adults with B-lymphoblastic leukemia (B-ALL) suffer relapse, and prognosis after relapse is very poor.^1,2^ Identifying those at risk for treatment failure and devising novel approaches to mitigate their relapse risk is imperative to improving outcome.

In B-ALL, deletions and mutations of the IKAROS family zinc finger 1 (*IKZF1*) gene are associated with an increased risk of relapse.^3^ *IKZF1* encodes the protein IKAROS which is a master lymphoid regulatory transcription factor^4,5^ and chromatin remodeler.^6^ One of the earliest regulators of lymphoid lineage identity,^7^ IKAROS is required for terminal B-cell maturation,^8,9^ and its loss of function is an important factor in differentiation-arrested, immature B-cell leukemias.

Various types of lesions in *IKZF1* may occur in B-ALL patients, including mono- or biallelic deletion of the entire gene, intragenic deletions, and loss of function mutations.^10^ The most common intragenic deletion (found in ~23% of adult and pediatric B-ALL) results in exclusive expression of the short IK6 isoform^11,12^ secondary to deletion of a 50 kilobase (kb) intragenic region^13^ (containing exons 4-7), and the resulting loss of four critical DNA-binding zinc fingers.^14^ IK6 retains the capacity to dimerize with residual wild-type IKAROS, inhibiting its function, thus IK6 is a dominant-negative inhibitor of IKAROS function. When retrovirally overexpressed, IK6 result in widespread transcriptional changes that promote cancerous proliferation, as cells are able to bypass typical metabolic restrictions imposed by tumor suppressive transcriptional regulators.^15^ Additionally, studies using inducible IK6 mice have shown the IK6 isoform also promotes carcinogenesis by mediating the transition to stromal independence via disruption of integrin signaling pathways.^16^

While *IKZF1* mutations and deletions are clinically correlated with poor outcome and increased risk of relapse, indicating *IKZF1* status could be a useful prognostic marker, clinical trials to modulate therapeutic intensity based on *IKZF1* status have had mixed results.^17–19^ How the broad spectrum of *IKZF1* genetic variants seen in B-ALL affect chemosensitivity and outcome and whether *IKZF1* status is a relevant prognostic criterion only in the context of certain driver translocations/mutations remain unclear.^20^

Some previously published in vitro studies using RNAi-mediated knockdown of *IKZF1* have shown evidence of resistance to corticosteroids,^11,21^ but others report no change in chemosensitivity.^22^ These mixed results may be in part due to variable reduction in IKAROS protein with RNAi strategies. Additionally, previous studies on the impact of the IK6 isoform in B-ALL have been limited by reliance on viral overexpression^14,15,23,24^ that may not accurately model the relative expression level of IK6 to other *IKZF1* isoforms. To overcome the limitations of these previous model systems, we used CRISPR/Cas9 strategies to generate a series of human B-ALL cell lines with *IKZF1* lesions mirroring those found in patients. We generated isogenic cell lines heterozygous or null for *IKZF1* by targeting early exons with efficient single guide RNAs (sgRNA). We also developed a CRISPR/Cas9 homology directed repair (HDR) strategy in which the IK6 isoform is knocked-in to the endogenous locus. This system allows for the expression of IK6 under the complete control of the endogenous promoter while simultaneously removing the expression of one wildtype allele, genetically recapitulating the heterozygous IK6-generating intragenic deletion occurring in B-ALL patients. With this series of human B-ALL cell lines, we investigated the genotype-specific differences in multi-drug resistance, gene expression profiles, in vivo engraftment, and cell cycle kinetics.

## Methods

### Animal models

Animal procedures were approved by the Institutional Animal Care and Use Committee (IACUC) of Baylor College of Medicine. NOD *scid* gamma mice (NSG; The Jackson Laboratory #005557) were housed in a dedicated immunocompromised suite. For xenograft transplants, leukemia cell lines were washed twice with sterile PBS, resuspended at a concentration of 1×10^6^ cells/100μL in sterile PBS, and injected retro-orbitally under brief isoflurane anesthesia. Periodic retro-orbital blood samples were taken by capillary tube from the eye contralateral to the leukemia cell injections. Animals were monitored and euthanized when moribund according to standard IACUC procedural guidelines to minimize pain and distress.

### Generation of isogeneic human B-ALL cell lines with *IKZF1* lesions

Using CRISPRscan (http://www.crisprscan.org),^25^ we selected a series of 6 sgRNAs targeting exon 2 (the first coding exon) or exon 3 of *IKZF1*. sgRNAs were chosen to minimize predicted off-target effects with maximum cleavage efficiency and were synthesized by in vitro transcription as previously described^26^ using the HiScribe T7 High Yield RNA Synthesis Kit (New England Biolabs, E2040S) or, alternatively, were commercially synthesized with 2’-O-methyl 3’ phosphorothioate modifications in the first and last three nucleotides (Synthego). We introduced the CRISPR/Cas9 system by electroporation with sgRNA-Cas9 ribonucleoprotein complexes (RNPs) to generate *IKZF1* mutant clones as previously described^26^ using the Invitrogen Neon Transfection System (10μL tip, 1400V, 35ms single pulse). We identified single cell-derived clonal lines with *IKZF1* frameshift mutations in one or both alleles by Sanger sequencing and the TIDE decomposition algorithm prediction (https://tide.nki.nl).^27^ Ablation of protein expression was confirmed by immunoblotting. Custom double-stranded DNA homology-directed repair (HDR) templates were synthesized by Twist Biosciences (see Supplemental Methods for complete sequences). Clonal knock-in cell lines were generated by the same method with the addition of 500ng HDR template DNA to the sgRNA-Cas9 RNPs as previously described.^28^ For both the IK6-GFP fusion tag and the lentiviral GFP transduced cell lines, GFP-positive cells were sorted on a BD FACS Aria III cytometer. Propidium iodide was used as dead cell exclusion stain. The plenti-CAG-IKZF1-V1-FLAG-IRES-GFP plasmid used in the rescue experiments was a gift from William Kaelin (Addgene plasmid # 107387).^29^

### RNAseq and qRT-PCR analysis

RNA was isolated from clonal leukemia cell lines using the RNeasy Plus Mini kit (Qiagen, 74134). RNAseq libraries were prepared using the TruSeq Stranded mRNA Library Prep (Illumina, 20020594) and the sequencing was done on the Illumina NextSeq 500 instrument (single end 75 bp reads). Single-end RNAseq reads were mapped to human genome (hg19) using Hisat 2 version 2.1.0 with default parameters. Aligned reads were then converted to BAM files and indexed using samtools version 1.3. Reads were quantified and then normalized using cuffquant and cuffnorm from the Cufflinks pipeline (version 2.2.1). Reads mapped to non-coding regions were masked from analysis. Differential gene expression was performed using R (3.6.2) on FPKM normalized tables using fold change ≥2. For pairwise comparisons, statistical significance was calculated using a pairwise t-test, and Bonferroni-corrected values were utilized in cases with multiple comparisons.

cDNA was made using the iScript cDNA Synthesis Kit (BioRad, 170-8891) with 1μg of RNA. Taqman qPCR was performed using the Luna^®^ Universal qPCR Master Mix (New England Biolabs, M3003) using the C1000 Touch Thermal Cycler (BioRad). All FAM-labeled Taqman assay probes were purchased from Thermo Fisher (Applied Biosystems) (product information in the Supplemental Methods).

### Microscopy

Fluorescence imaging was done by confocal microscopy of live cells in a glass culture dish using Hoechst 33258 as a nuclear stain. All images were taken on the GE Healthcare DeltaVision LIVE High Resolution Deconvolution Microscope using an Olympus 40X U Apo/1.35 NA oil objective with iris and analyzed with SoftWoRx.

### Statistical Analysis

Statistical comparisons were evaluated using the Student’s t-test or one-way ANOVA as indicated using GraphPad Prism 7. All data are presented as mean ± SEM. For animal survival studies, Kaplan-Meier survival curves show time to morbidity and were analyzed using the logrank test. Where appropriate, exact calculated p-values are shown. n indicates the number of biological replicates within each group.

Additional information is available online in the accompanying Supplemental Methods.

## Results

### Electroporation delivery of Cas9/sgRNA RNPs efficiently ablates IKAROS expression in human B-ALL cell lines

In order to identify critical transcriptional network alterations that could relate to changes in cell growth, signaling, and chemotherapy response, we engineered a set of clonal cell lines with accurately determined reduction or complete ablation of IKAROS protein expression. First, we electroporated the Nalm-6 B-ALL cell line with recombinant Cas9 protein precomplexed with sgRNAs (RNPs) targeting the first coding exons of *IKZF1* (Figure 1A). To ensure our electroporation conditions were optimized, as a control we used an sgRNA against *CXCR4* which encodes a cell surface adhesion protein. After 72 hours, 59.2% of Nalm-6 cells electroporated with Cas9-*CXCR4* sgRNA RNP showed ablation of cell surface expression of CXCR4, confirming efficient delivery of Cas9/sgRNA (Figure 1B). Sanger sequencing of DNA from the electroporated cell pools of PCR amplicons surrounding the *IKZF1* target sites confirmed the high occurrence of indels, particularly for 3 of the 6 sgRNAs tested (sg55, 57 and 79) and all further experiments were done with these 3 sgRNAs.

**Figure 1:**
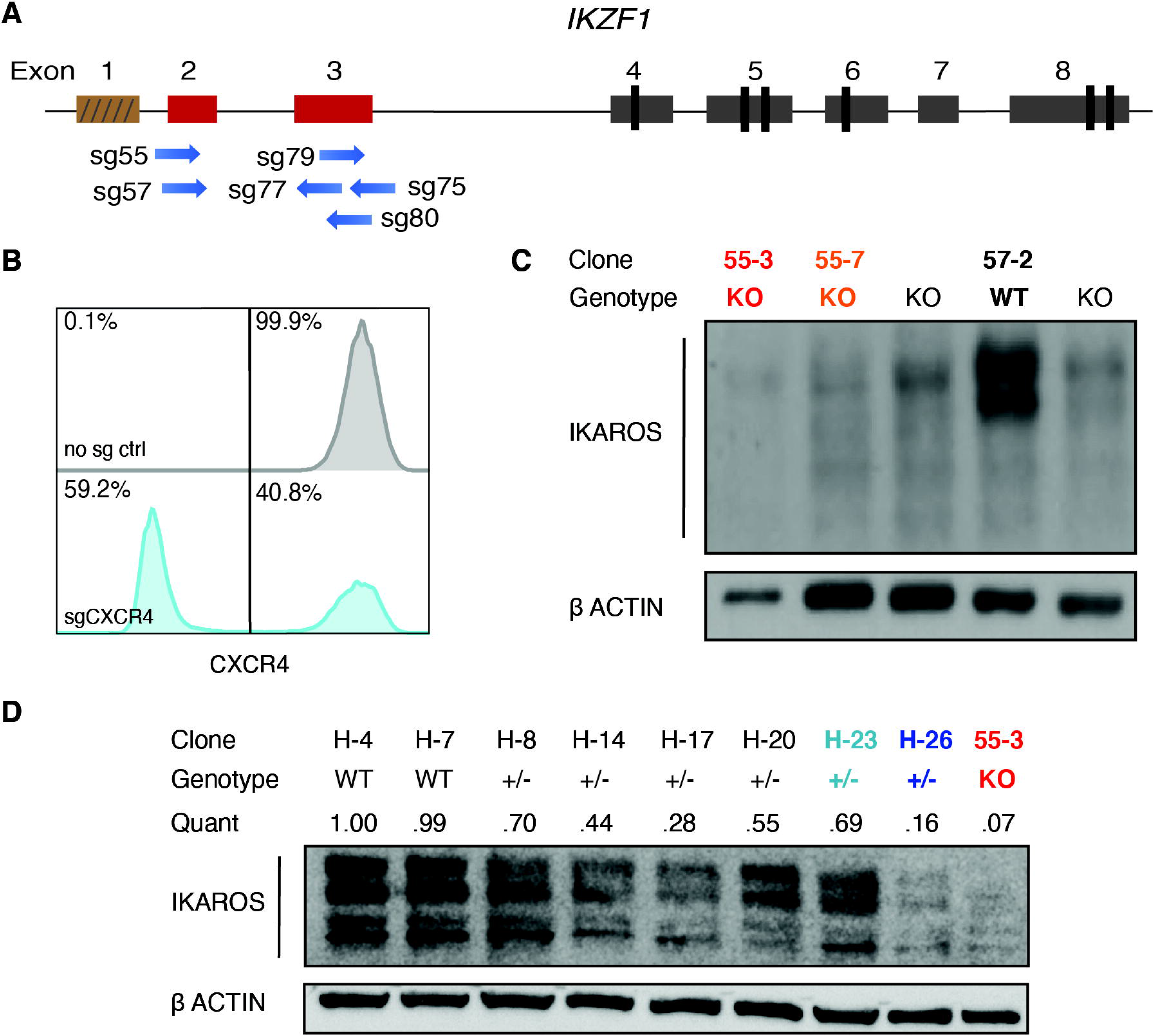
Cas9-sgRNA RNP targeting of *IKZF1* efficiently eliminates protein expression in Nalm-6 cells. A) sgRNAs targeting the early coding exons of *IKZF1* (exons 2 and 3, shown in red; brown indicates non-coding exon 1). B) Flow cytometry histograms showing CXCR4 cell-surface protein using sgRNA against CXCR4 as a control marker for CRISPR efficiency in Nalm-6. C) IKAROS immunoblotting of single-cell derived Nalm-6 clones with Sanger sequencing-verified *IKZF1* frameshift mutations. D) Reduced concentration RNP delivery yields a higher frequency of heterozygous clones with varying protein levels. Relative quantification was normalized to ACTIN. Bold font/coloring of clones on C) and D) indicate those used in further experiments that are similarly color-coded throughout all figures.

We derived and sequenced clonal cell lines and inferred their suspected indels from TIDE analysis. For subsequent experiments, we selected clonal lines with complete ablation of IKAROS protein expression (Figure 1C), which are referred to throughout as *IKZF1* KO. Our initial approach generated no predicted heterozygous samples; 74% of edited clones sequenced had indels on both alleles (data not shown). In order to engineer clones with monoallelic *IKZF1* lesions, we decreased the amount of Cas9 protein and sgRNA by 50% and 75% and successfully isolated clones with monoallelic, out-of-frame indels from these conditions with reduced IKAROS protein expression ranging from 16-70% of WT expression levels (Figure 1D), hereafter referred to as *IKZF1* +/-. As controls, we selected clonal lines with no evidence of *IKZF1* editing by Sanger sequencing and expected levels of IKAROS protein expression, referred to throughout the text as *IKZF1* wildtype (WT). IKAROS protein is known to regulate its own expression level,^30^ and therefore, as expected, knockout clones showed a 3- to 4-fold upregulation of frameshift mutant *IKZF1* mRNA by qRT-PCR (Supp. Figure S1).

We also created a similar panel of heterozygous and knockout clones using the Tanoue B-ALL cell line and confirmed loss of IKAROS protein (Supp. Figure S2A).

### *IKZF1* deletion results in a stem-like expression signature and upregulates targetable pathways

In order to identify the spectrum of gene expression changes in response to loss of IKAROS, we performed RNAseq on our clonal cell lines. Gene set enrichment analysis (GSEA) of the resulting significantly up- or down-regulated genes in KO clones showed a strong enrichment of a hematopoietic stem cell (HSC) expression signature (consistent with previously published murine models),^11^ and an acute myeloid leukemia (AML) stem cell expression signature (Figure 2A). Additionally, genes associated with the IL-7/JAK/STAT pathway were significantly altered between WT and KO cells (Figure 2A), consistent with reported trends in expression profiling of ALL patient samples.^12^ We also used the RNAseq reads spanning *IKZF1* exon 2 to confirm the CRISPR/Cas9 induced in/dels (Supp. Figure S3). We selected a subset of previously published IKAROS target genes^31^ for validation of the RNAseq results by qRT-PCR and found similar trends in gene expression changes using both methods (Supp. Figure S4).

**Figure 2:**
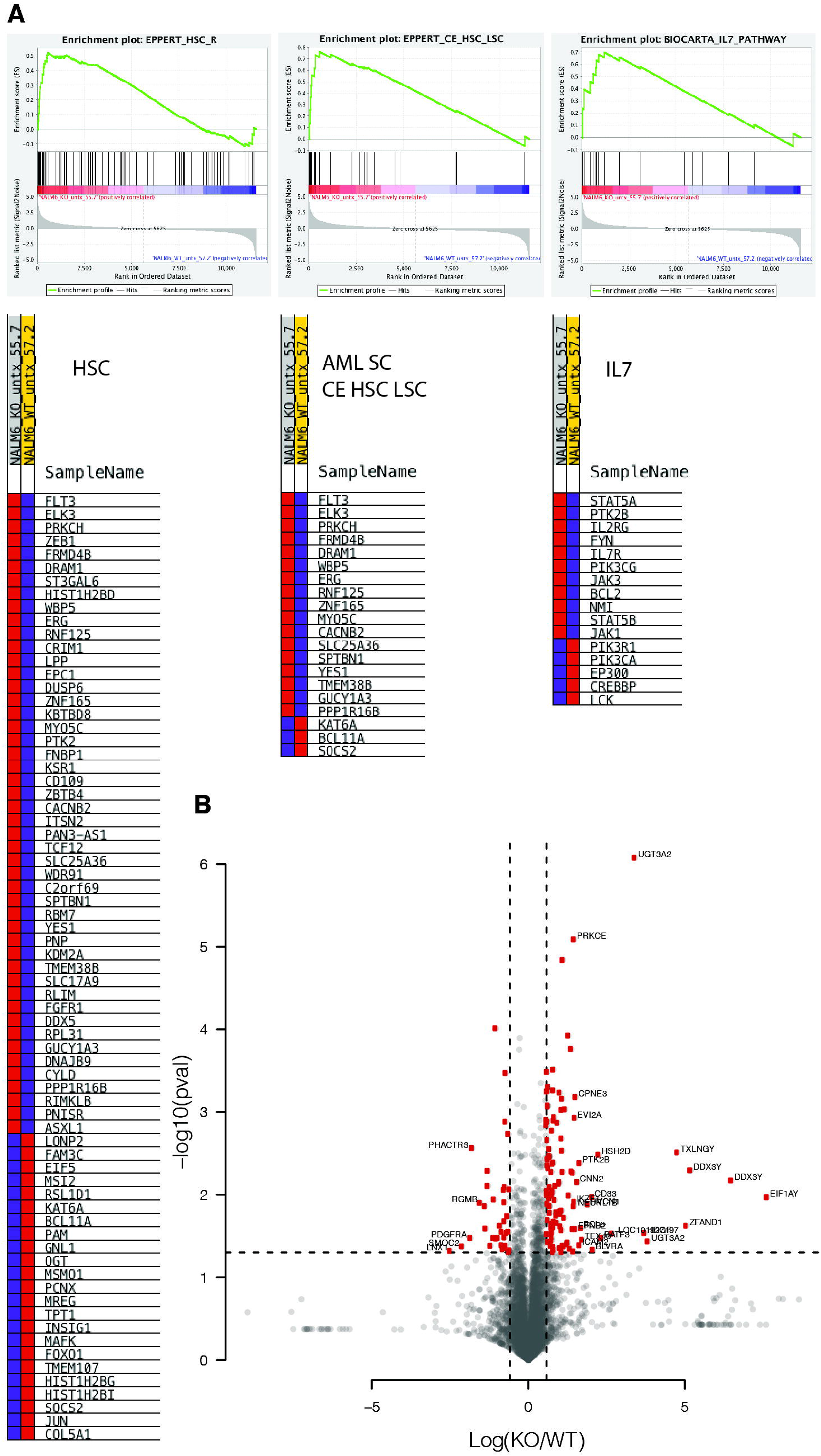
RNAseq analysis after IKAROS depletion reveals a stem cell-like gene expression pattern and upregulation of acute myeloid leukemia stem cell-associated genes. A) Gene set enrichment analysis (GSEA) shows gene expression changes in signatures associated with hematopoietic stem cells (HSC), AML leukemia stem cells, (AML SC CE HSC), and the IL7/JAK/STAT pathways after *IKZF1* knockout. B) Volcano plot showing the most statistically significantly up- and down-regulated genes (*IKZF1* KO vs. WT).

We noted a number of the genes most down-regulated after *IKZF1* KO (e.g., *PDGFR*) are involved in regulation of various cell cycle-dependent processes (Figure 2B) so we performed cell cycle analysis in our edited Nalm-6 cells, finding a higher proportion of KO cells in G1 phase in untreated samples (Supp. Figure S5A). These results suggested a relatively quiescent state, which aligns with the identified stem cell-like gene expression signature. WST-1 assays also revealed a substantially lower baseline metabolic rate in KO cells (Supp. Figure S5B). To track the relative growth rate of the B-ALL cells, we transduced our isogenic lines with a lentiviral GFP expression system and conducted an in vitro competition assay with WT-GFP versus KO and the WT gradually out-grew the KO over 10-12 days (Supp. Figure S5C-D). These in vitro data suggest that *IKZF1* deletion promotes not only a stem cell-like gene signature but also a stem cell-like phenotype.

### *IKZF1* deletion enhances bone marrow engraftment and shortens survival in mouse xenograft models

To determine the impact of IKAROS loss on bone marrow homing and engraftment of human BALL, we transplanted NSG mice intravenously with either the *IKZF1* KO or WT Nalm-6 clones. Peripheral blood sampling on day 18, before noticeable signs of disease, showed a significantly higher engraftment in the animals transplanted with *IKZF1* KO cells compared to WT (Figure 3A). Mice harboring *IKZF1* KO B-ALL cells succumbed to leukemia significantly earlier than the WT group (Figure 3B). At the time of sacrifice, *IKZF1* KO xenografted animals showed trends toward higher engraftment compared to WT in bone marrow and spleen, but a statistically significant difference was only observed in liver (Figure 3C). Similar results were seen with *IKZF1*-deleted Tanoue B-ALL cell xenografts (Supp. Figure S2B). These results together suggest that deletion of *IKZF1* leads to more efficient motility and homing of B-ALL to the bone marrow and a more rapidly progressing, infiltrative disease.

**Figure 3:**
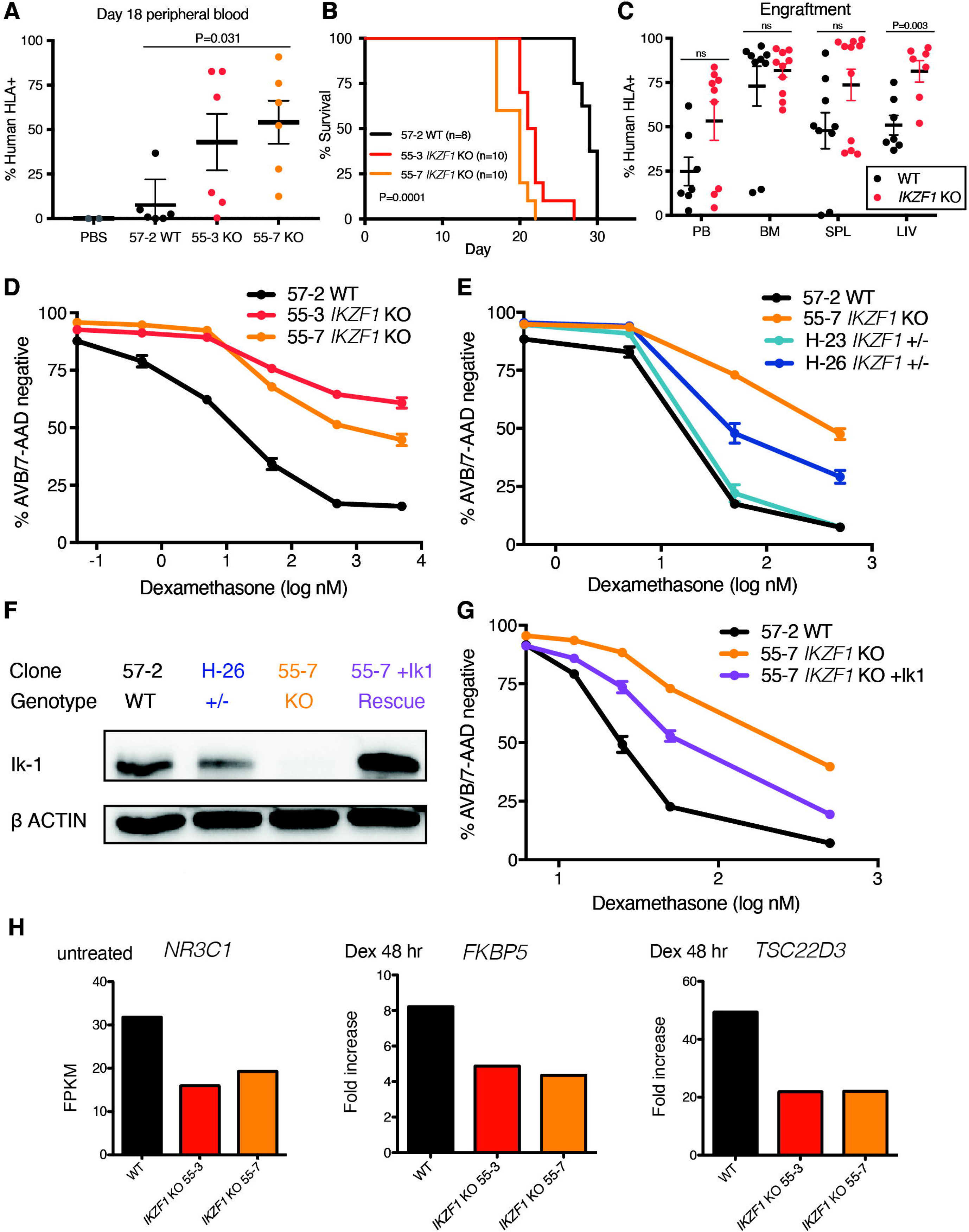
*IKZF1* deletion reduces survival in NSG mice and leads to profound dexamethasone resistance that is rescuable by IK1 overexpression. (A-C) NSG mice xenografts transplanted with Nalm-6 clonal cell lines: A) Flow cytometry for human HLA-ABC marker in peripheral blood samples taken day 18. B) Kaplan-Meier survival curve, n= total number of mice in each group. C) Tissue engraftment of Nalm-6 cells at time of sacrifice; PB= peripheral blood, BM= bone marrow, SPL= spleen, LIV= liver. D), E), & G) 72-hour dexamethasone treatment of Nalm-6 clonal cell lines showing percent AnnexinV/7-AAD double negative population as measured by flow cytometry. Samples were done in triplicate, error bars show SEM. F) IKAROS immunoblotting of single-cell derived Nalm-6 clones and IK1 lentivirus overexpression (55-7+ IK1 rescue shown in purple). G) shows partial restoration of dexamethasone sensitivity in *IKZF1* KO after IK1 rescue. H) RNAseq gene expression levels plotted as Fragments per Kilobase of Transcript per million mapped reads (FPKM) in untreated samples (*NR3C1*) or fold increase of FPKM after 48 hour, 50nM dexamethasone treatment (*FKBP5* and *TSC22D3*).

### *IKZF1* deletion confers B-ALL with profound resistance to glucocorticoids (GC) by blunting response of GC receptor target genes

We treated our *IKZF1* KO and WT Nalm-6 clonal cell lines with increasing doses of the GC dexamethasone. While the *IKZF1* WT cells underwent a dose- and time-dependent inhibition of proliferation and induction of apoptosis, the *IKZF1* KO cells were highly resistant to these GC-induced effects (Figure 3D and Supp. Figure S6A). The heterozygous clone (H-26; ~16% protein expression compared to WT, Figure 1D) displayed an intermediate resistance (Figure 3E), consistent with previously published findings using RNAi.^31^ Another heterozygous clone (H-23) with ~69% of WT protein expression level (Figure 1D), showed no change in resistance (Figure 3E). We observed similar results in the clonal Tanoue cell lines (Supp. Figure S2C).

In order to substantiate the claim that the observed drug resistance phenotypes are a direct result of *IKZF1* depletion, we introduced the full-length protein isoform IK1 using a bicistronic GFP lentiviral overexpression system into the 55-7 Nalm-6 *IKZF1* KO cell line. After sorting for GFP-positive cells, we confirmed IK1 overexpression (Figure 3F). When treated with dexamethasone we observed a partial rescue of glucocorticoid sensitivity in the IK1 overexpressing cells (Figure 3G), confirming IKAROS depletion as the driver of resistance.

We also performed RNAseq on Nalm-6 WT and *IKZF1* KO clones after dexamethasone treatment. We selected the 50nM dose at both the 24-hour and 48-hour timepoints because there was a small but detectable difference in apoptosis between WT and KO, while minimizing bias from inclusion of RNA from apoptotic cells (Supp. Figure S6B).^32^ In the KO cells we observed lower baseline expression of *NR3C1*, the gene encoding the GC receptor, and a blunted response of several downstream GC receptor transcriptional target genes including *FKBP5* (a prolyl isomerase modulator of GC receptor sensitivity) and *TSC22D3* (an anti-inflammatory, GC-induced leucine zipper) (Figure 3H). These studies provide direct evidence for *IKZF1* KO-driven glucocorticoid resistance.

### *IKZF1* modulation alters sensitivity to chemotherapies in vitro

To test whether the observed effect was unique to glucocorticoids, we examined the impact of loss of IKAROS expression on sensitivity to a variety of chemotherapeutic agents used for the treatment of B-ALL. We found that full and partial ablation of IKAROS in Nalm-6 resulted in relative resistance to vincristine (Figure 4A and Supp. Figure S7A), daunorubicin (Figure 4B), and asparaginase (Figure 4C). We found no difference in sensitivity to methotrexate between the *IKZF1* KO and WT cells (Figure 4D). The *IKZF1*-KO Tanoue cells similarly displayed moderate resistance to vincristine and asparaginase (Supp. Figure S2D-E).

**Figure 4:**
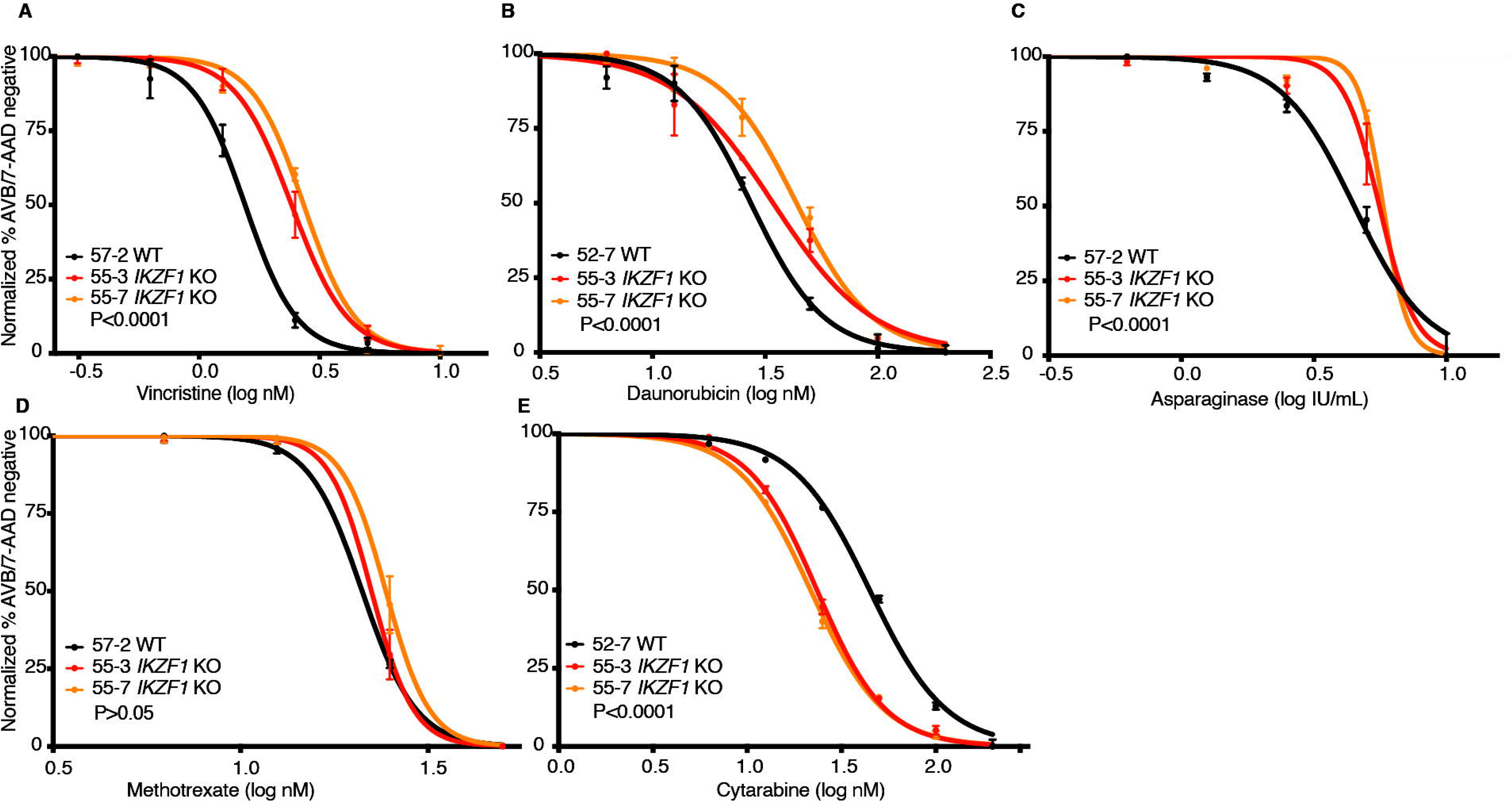
Deletion of *IKZF1* leads to broadly increased chemoresistance. Drug treatment of Nalm-6 clonal cell lines showing percent AnnexinV/7-AAD double negative population as measured by flow cytometry. Samples were done in triplicate, error bars show SEM. All assays were 72 hours except A) vincristine, 48 hours. B) daunorubicin, C) asparaginase, D) methotrexate (no statistically significant difference), and E) cytarabine.

Interestingly, when treated with the nucleoside analog cytarabine, we found that the *IKZF1* KO and heterozygous cells were more sensitive than WT (Figure 4E and Supp. Figure S7B; Supp. Table S1 contains a summary of all IC_50_ values), suggesting nucleoside analogs may be a particularly effective class of drugs for *IKZF1*-deleted B-ALL.

### GFP-tagged IK6 knock-in to the endogenous locus mimics the intragenic deletion found in patients

While most clinical B-ALL studies group all *IKFZ1* lesions together, some have found that different lesions have distinct effects on outcome. In particular, studies examining the prognostic impact of the IK6-generating intragenic deletion of exons 4-7 have been conflicting. ^17–19,33–37^ Further, in addition to inhibition of residual wild-type IKAROS, the IK6 isoform also heterodimerizes with other IKAROS family proteins such as HELIOS (encoded by *IKZF2*) and AIOLOS (encoded by *IKZF3*), inhibiting their function. Therefore, the transcriptomic and phenotypic impact of IK6 could differ from deletion of the entire *IKZF1* gene. Thus, we expanded our panel of *IKZF1* alterations with the creation of a novel IK6 dominant-negative isoform expressed under the control of the endogenous *IKZF1* locus.

We employed an HDR strategy in which exon 3 was directly fused to exon 8 on a 3.2kb linear dsDNA template, with an eGFP fusion tag on the c-terminus, serving the dual purpose of selection by cell sorting and tracking of intracellular localization (Figure 5A). This strategy successfully generated GFP-expressing cells (efficiencies of 0.1-0.4% of live cells) (Figure 5B). We sorted GFP-positive cells, one cell per well, into 96-well plates. After expansion, we selected clones based on PCR and Sanger sequencing that had one wildtype unedited allele with the other allele showing perfect homologous recombination matching the template sequence.

**Figure 5:**
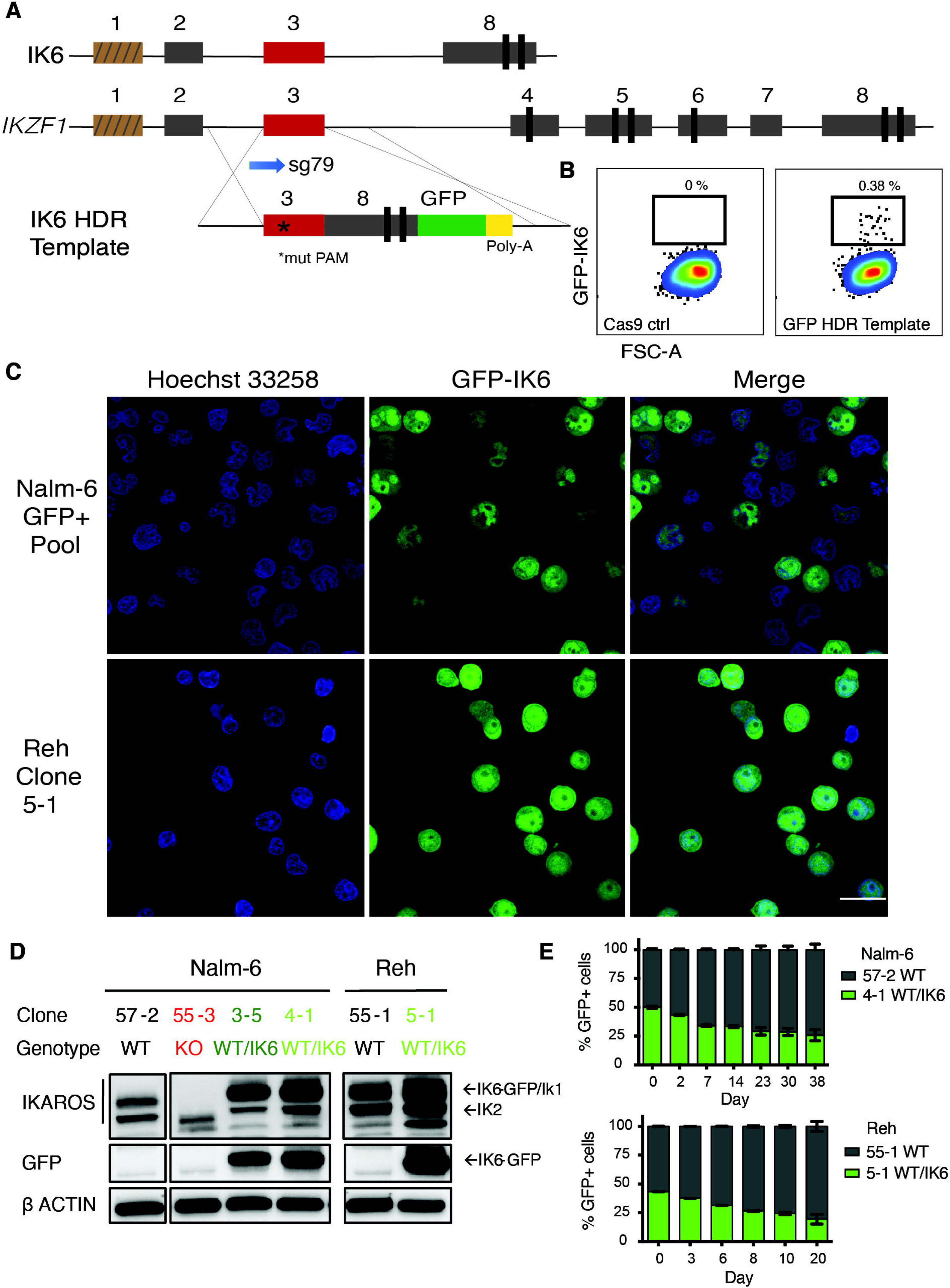
Creation and characterization IK6-expressing B-ALL via knock-in of IK6 into the endogenous *IKZF1* locus. A) Homology-directed repair strategy for GFP-fused IK6 knock-in using 3.2kb dsDNA HDR template. (targeted exon 3 shown in red; brown indicates non-coding exon 1; *mutated protospacer-adjacent motif or PAM to prevent cleavage of template). B) Flow cytometry plots showing GFP+ sort gate used to purify cells that had IK6-GFP knock-in. C) IK6-GFP fusion protein in nucleus and also mislocalization to cytoplasm as shown by live cell confocal microscopy. Upper row, Nalm-6 GFP-positive pool showing heterogenous expression one month after sorting and lower row, Reh single cell-derived clone. All images were taken on the GE Healthcare DeltaVision LIVE High Resolution Deconvolution Microscope using an Olympus 40X U Apo/1.35 NA oil objective with iris and analyzed with SoftWoRx. Scale bars are 20μm. D) Immunoblotting for IKAROS and GFP-IK6 fusion protein in Nalm-6 and Reh single-cell derived clones. E) In vitro competition of Nalm-6 (upper graph) and Reh (lower graph). GFP was measured by flow cytometry at indicated timepoints. All samples were done in triplicate, error bars show SEM.

We observed the IK6-GFP fusion protein, as expected,^38^ partially aberrantly localized to the cytoplasm (Figure 5C). Immunoblot of whole cell extracts showed IK6 was more highly expressed than the normal IK1/IK2 isoforms found in the wildtype clone (Figure 5D). This phenomenon is commonly seen in cell lines and patient samples expressing IK6, presumably owing in part to reduced degradation secondary to lack of a ubiquitin ligase-interacting domain.^39^ Both the Nalm-6 and Reh WT/IK6 cells proliferated less than WT over several weeks of coculture, consistent with the *IKZF1* KO cell lines and suggestive of a relatively quiescent, stem cell-like phenotype (Figure 5E). Thus, with our CRISPR/Cas9 HDR strategy we successfully generated B-ALL cells with expected IK6 expression and cellular localization, ideal for direct comparison to *IKZF1* WT, KO and +/- to precisely delineate the impact of these various *IKZF1* genotypes commonly encountered in human B-ALL.

### Impact of *IKZF1* genotype on gene expression

To assess gene expression changes induced by the different *IKZF1* genotypes, we performed RNAseq on our engineered Nalm-6 cell lines, comparing *IKZF1* WT, KO, +/- and IK6-GFP/WT cells. Isogenic lines of the same *IKZF1* genotype clustered together (Figure 6A). In general, the differentially expressed genes were similarly expressed across all the *IKZF1* mutated lines, with the most pronounced effects in most genes noted in the *IKZF1* KO (Figure 6A, B). We validated the findings at the protein level by flow cytometry of several clinically relevant, cell surface-expressed phenotyping markers including CD19, CD22, CD33, and FLT3 (Figure 6C). This indicates that in B-ALL *IKZF1* lesions result in a similar gene expression pattern, with the magnitude of gene expression changes influenced by the amount of residual IKAROS function.

**Figure 6:**
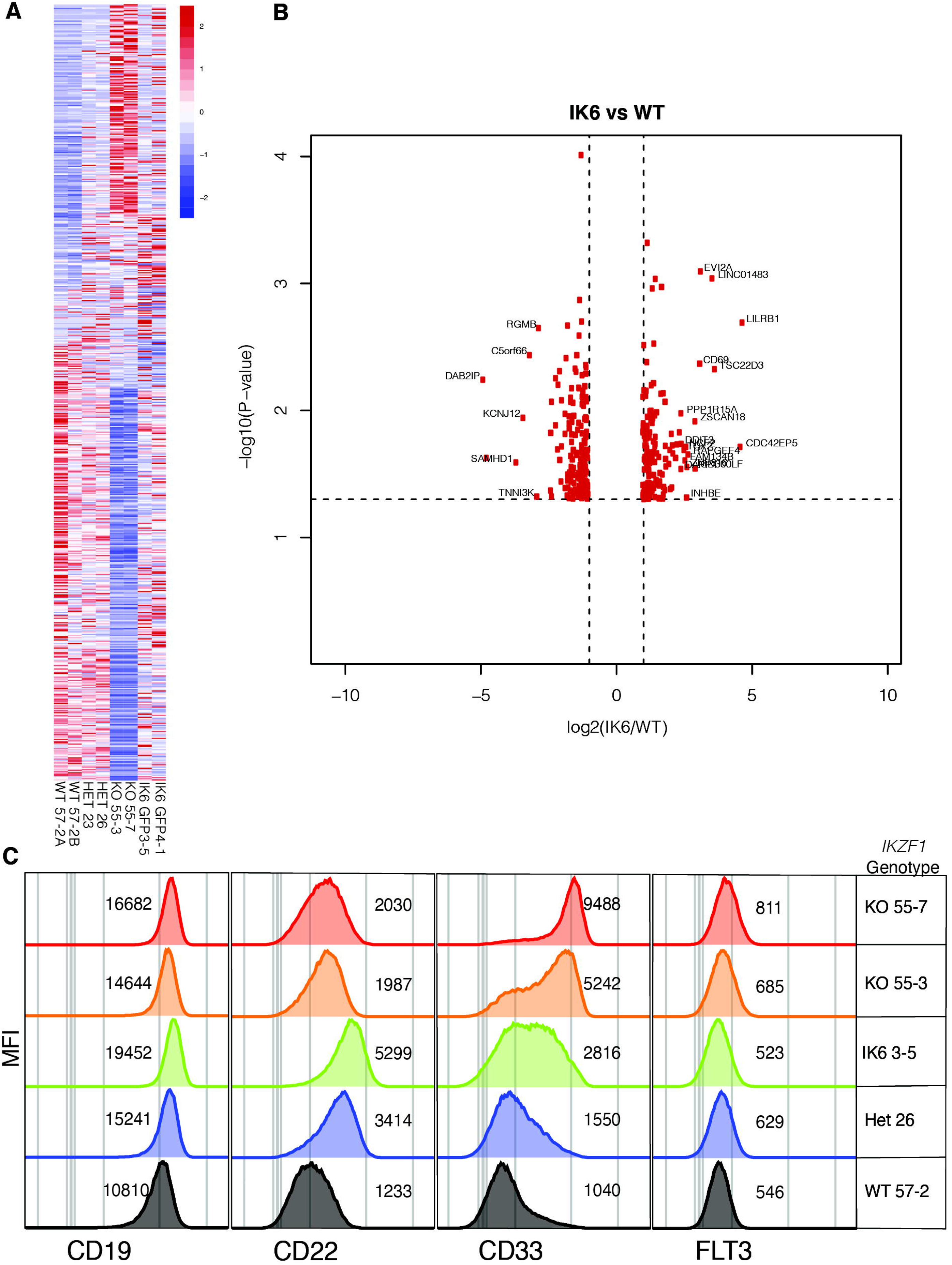
IK6 knock-in shows similar changes in gene expression patterns to heterozygous knockout. All samples are from Nalm-6 clonal cell lines. A) Heatmap showing differential gene expression analysis from RNAseq from wild type (WT), heterozygous (HET), knockout (KO) and IK6 knockin. B) Volcano plot highlighting the most significantly up- or down-regulated genes (IK6 vs. WT). C) Flow cytometry histograms showing confirmation of protein expression changes of a panel of cell surface markers commonly used in leukemia phenotyping. MFI, mean fluorescence intensity.

### IK6 knock-in transplanted NSG mice have accelerated disease progression compared to WT

To establish the impact of IK6 expression on bone marrow homing and engraftment, we repeated the NSG mouse xenograft experiment with the Nalm-6 WT/IK6 cells compared to WT and KO counterparts. Peripheral blood engraftment of WT/IK6 Nalm-6 cells 2 weeks post-transplant was similar to WT, while KO engraftment was significantly higher as in our previous experiments (Figure 7A). Despite this lower early peripheral blood engraftment, mice transplanted with IK6 knock-in cells succumbed to the disease at approximately the same time as the KO transplanted animals (Figure 7B). Liver weight was significantly increased in the IK6 transplanted mice (Figure 7C) but spleen weight was not significantly different (Supp. Figure S8). These results suggest that, though IK6 was associated with increased liver infiltration with less early peripheral blood engraftment, IK6 and KO contribute similarly to overall B-ALL disease progression.

**Figure 7:**
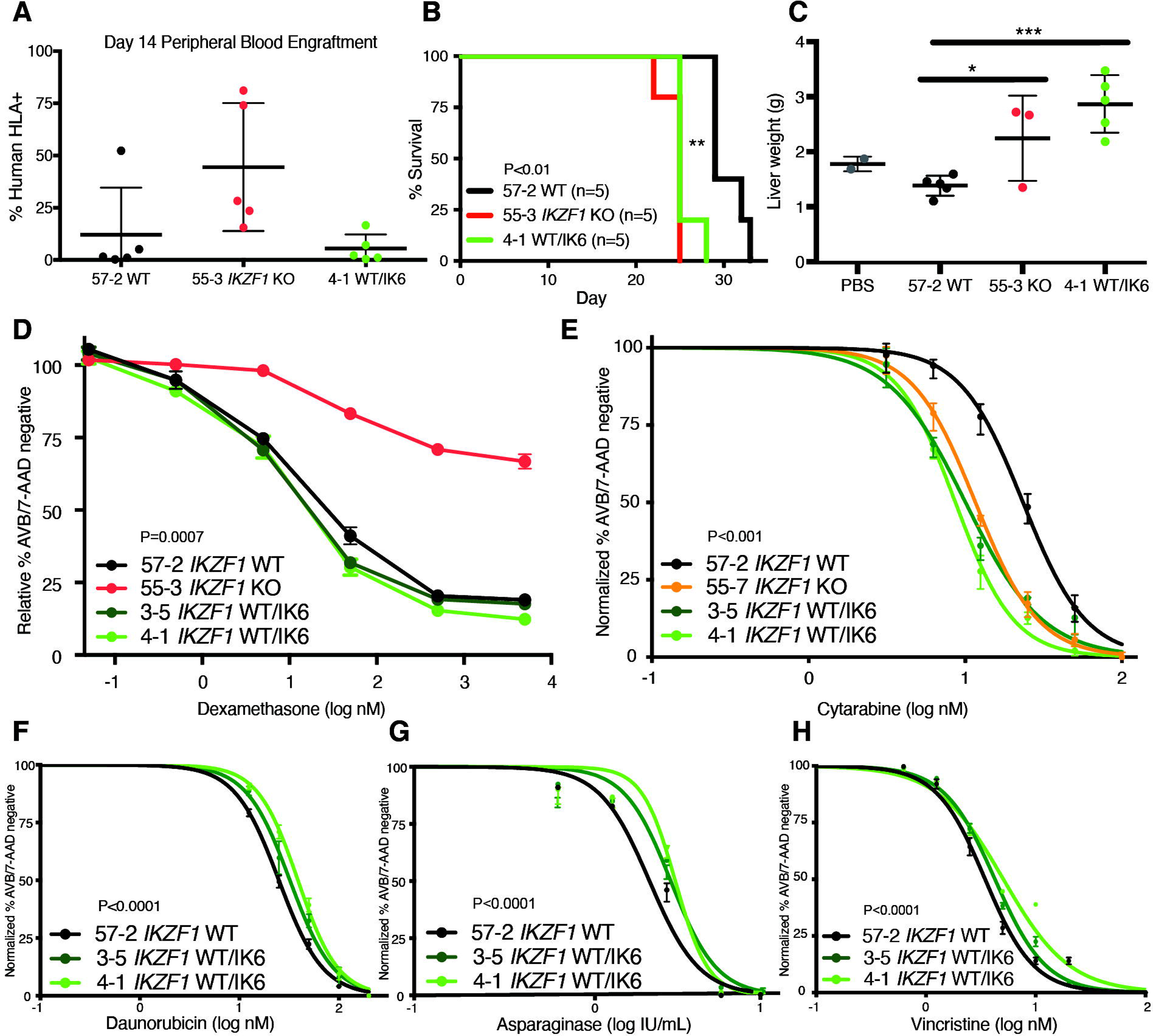
IK6 knock-in leads to shortened survival time in NSG mice with increased extramedullary organ engraftment. NSG mice xenografts transplanted with Nalm-6 clonal cell lines (A-C): A) Flow cytometry for human HLA-ABC marker in peripheral blood samples taken day 14. B) Kaplan-Meier survival curve, n= total number of mice in each group. C) mouse liver weights at time of sacrifice. (D-H) drug treatment of Nalm-6 clonal cell lines showing percent AnnexinV/7-AAD double negative population as measured by flow cytometry after 72 hours of treatment with D) dexamethasone, E) cytarabine, F) daunorubicin, G) asparaginase, and H) after 48 hours of treatment with vincristine. Samples were done in triplicate, error bars show SEM.

### IK6 knock-in creates a unique drug resistance profile

To determine if B-ALL with IK6 expression had a different chemosensitivity profile compared to B-ALL with deletion of *IKZF1*, we treated WT and IK6 cell lines with chemotherapy. We observed no change in sensitivity to dexamethasone in IK6 cells compared to wildtype (Figure 7D). This data combined with data from our Nalm-6 *IKZF1*+/- clones with varying levels of residual IKAROS protein suggest that WT IKAROS protein levels must be severely reduced (to <20% normal levels) in Nalm-6 cells to observe changes in resistance to dexamethasone. Conversely, the WT/IK6 cells displayed moderate resistance to daunorubicin, asparaginase, and vincristine, similar to the *IKZF1* KO and *IKZF1*+/- clones (Figure 7F-H and Supp. Table S1 containing summary table of IC_50_ values). Also, like the *IKZF1* KO and *IKZF1* +/-, the WT/IK6 cells demonstrated increased sensitivity to cytarabine (Figure 7E). Overall, these studies suggest that considering the specific type of *IKZF1* lesions may inform treatment decisions.

## Discussion

In order to detect small changes in chemoresistance that would have been previously undetectable using knockdown approaches, we generated complete IKAROS protein knockout in human cells. RNAseq on *IKZF1* knockout in B-ALL cell lines revealed a stem cell-like gene expression pattern, with relative quiescence, enhanced bone marrow engraftment, and resistance to most chemotherapeutic agents. These results are consistent with the relapsing phenotype observed in patients with *IKZF1* deletions, which are associated with increased risk of relapse in clinical studies. To date, some cancer consortia have incorporated *IKZF1* status along with bone marrow minimal residual disease (MRD) into their risk-stratification schema, defining patients with *IKZF1* mutations/deletions and MRD positive disease as high risk and assigning them to intensified post-induction therapy.^40^ Other published studies using varying methods of measurement and statistical analyses have shown, however, little benefit of *IKZF1* status as a classifier in risk stratification,^35^ though these data are from a relatively small cohort (reviewed in Olsson and Johansson^41^). Overall, our studies demonstrate that lesions of *IKZF1* result in cell-intrinsic resistance to most ALL-directed chemotherapy.

We identified patterns of chemotherapy sensitivity in *IKZF1* mutant B-ALL that may have important clinical implications. Complete ablation of IKAROS protein expression in BALL cells resulted in profound dexamethasone resistance, while a different pattern was observed with other genotypes. In the sequencing-confirmed *IKZF1*+/- isogeneic clones, we found that IKAROS protein expression varied, which allowed us to determine relative corticosteroid resistance on the basis of protein level. Interestingly, we only observed a measurable effect on dexamethasone sensitivity in clones with very low residual IKAROS levels (<20%). Additionally, while the chemosensitivity profile of the WT/IK6 clones is similar to that of the *IKZF1*-KO clones, the WT/IK6 cells maintained their sensitivity to dexamethasone. These finding suggest IKAROS is necessary for dexamethasone sensitivity with a threshold level of required protein expression, below which an IKAROS dose-dependent effect occurs. Additionally, while all *IKZF1* engineered lesions result in relative resistance to most standard ALL-directed agents, we found a uniform increased sensitivity to the nucleoside analog cytarabine. While low-dose cytarabine is included in a subset of B-ALL post-induction therapy cassettes, our findings suggest regimens with intensification of cytarabine may preferentially benefit patients with *IKZF1* deleted B-ALL.

Comparing the gene expression profile and cell surface protein expression of our engineered B-ALL cell lines with and without *IKZF1* deletion, we found a surprising upregulation of the myeloid-associated cell surface phenotypic markers CD33 and FLT3 with IKAROS ablation, which may provide additional targeted therapeutic vulnerabilities. The CD33-targeted monoclonal antibody therapy gemtuzumab ozogamicin has activity against CD33+ BALL in vitro, in patient-derived xenografts,^42^ and has successfully induced stable complete remission in patients.^43–45^ Additionally, we observed increased cell surface expression of CD19 and CD22, indicating deletions/mutations of *IKZF1* should not impact sensitivity to CD19- or CD22-targeted therapies such as blinatumomab, inotuzumab, or C19- and/or CD22-targeting chimeric antigen receptor T-cells (CAR-T). We also found upregulation of the potentially targetable JAK/STAT pathway with *IKZF1* loss. Importantly, *IKZF1* aberrations are highly prevalent in Ph+ and Ph-like B-ALL which are characterized by activation of signaling pathways due to gene fusions and/or mutations, including subsets with JAK/STAT-activating lesions. Notably, all our investigations were done in Ph-negative, non-Ph-like B-ALL cells. How *IKZF1* loss impacts signaling pathways in Ph+ and Ph-like leukemia will be important future questions.

Overall, our data support clinical studies reporting that *IKZF1*-mutated B-ALL is an aggressive, infiltrative, and treatment-resistant disease. Notable differences in drug response and *in vivo* dynamics in xenografts exist between *IKZF1* KO cells and IK6 knock-in cells. Detailed delineation of the exact *IKZF1* status in ALL patients at diagnosis may be informative in more accurately determining risk stratification and the most effective therapeutic regimen.

## Acknowledgements

The authors thank the members of the Goodell Laboratory for helpful discussions and Catherine Gillespie for review of the manuscript. The work was supported by the National Institutes of Health, National Cancer Institute K08 CA 201611 and K12 CA090433-11 (RER); Be The Light Charitable Foundation (RER); Golfers Against Cancer (RER); The Gillson-Longenbaugh Foundation (RER); Texas Children’s Hospital Department of Pediatrics Pilot Award (RER); Hyundai Hope on Wheels (RER); The Ladies Leukemia League (RER); American Society of Hematology HONORS Award (RG); and Alex’s Lemonade Stand (CJ). All flow cytometry experiments were performed at the Baylor College of Medicine Cytometry and Cell Sorting Core which is supported by the National Institutes of Health, National Cancer Institute P30 Cancer Center Support (grant CA125123) and the National Institutes of Health (grants RR024574 and S10 OD025251). Microscopy experiments were performed at the Baylor College of Medicine Integrated Microscopy Core which is supported by the National Institutes of Health, National Cancer Institute P30 Cancer Center Support (grant CA125123), the National Institutes of Health, National Institute of Diabetes and Digestive and Kidney Diseases (grant 56338-13/15), the Cancer Prevention and Research Institute of Texas (Cancer Prevention and Research Institute of Texas (grant RP150578), and the John S. Dunn Gulf Coast Consortium for Chemical Genomics. STR DNA fingerprinting was done at The University of Texas MD Anderson Cancer Center CCSG-funded Characterized Cell Line Core, with support from the National Institutes of Health, National Cancer Institute CA016672.

## Authorship Contributions

JHR, RG, AG, MCG, GM, TS, CJ, SB, CDDC, LB, and RER performed experiments; JHR, RG, MCG, LB, MAG, and RER designed experiments and analyzed experimental results; JMR and MCG performed bioinformatics analysis; JHR and RER wrote the manuscript; all authors reviewed and commented on the manuscript. The authors also gratefully acknowledge editorial assistance with the manuscript from Catherine Gillespie, Department of Cell and Gene Therapy, Baylor College of Medicine.

