## Supplemental Data for "Modeling *IKZF1* lesions in B-ALL reveals distinct chemosensitivity patterns and potential therapeutic vulnerabilities"

### Supplemental Methods

#### Cell line culture

The human leukemia cell lines Nalm-6 (CRL-3273) and Reh (CRL-8286) were purchased from ATCC. The Tanoue cell line (ACC 399) was purchased from DSMZ. The identity of all leukemia cell lines was verified by STR fingerprinting. Cells were cultured in RPMI-1640 media supplemented with 10% Fetal Bovine Serum (Corning), 100 IU/mL Penicillin, 100 µg/mL Streptomycin, and 1mM L-glutamine. Cells were kept in a humidified incubator at 37 °C with 5% $CO_2$ and regularly PCR tested for mycoplasma.

#### Flow cytometry and competition assay

GFP-positive and GFP-negative cells were plated at a 50/50 ratio on Day 0. Cells were passaged every 2-3 days and GFP positivity was determined at indicated timepoints by flow cytometry using propidium iodide as a dead cell exclusion stain. Samples were acquired on a Becton Dickenson LSR II flow cytometer and BD FACS DIVA software and data were analyzed using the FlowJo VX software package.

#### Immunoblotting

SDS-PAGE and immunoblotting were performed as previously described.[1] IKAROS antibody (5443S; 1:500), GFP-HRP conjugated antibody (2037; 1:1000) and anti-rabbit IgG HRP-conjugated antibody (7074S; 1:5000) were purchased from Cell Signaling Technology. GFP antibody (sc-9996; 1:1000), ß-actin HRP-conjugated antibody (sc-47778; 1:2000), and anti-mouse IgG HRP-conjugated antibody (sc-516102; 1:5000) were purchased from Santa Cruz Biotechnology, Inc. All blocking and antibody incubations were done with 1X Tris-Buffered

Saline (TBS; pH 7.4) with 0.1% Tween and 5% bovine serum albumin. Images were collected using a Biorad Chemidoc MP imaging system.

**Annexin V/7-AAD assay**

Cells were cultured in 96-well plates at a concentration of 200,000 cells/mL for 48-72 hours with increasing doses of indicated drugs or vehicle control. 7-AAD (559925) and Annexin V APC (550475) were purchased from BD Biosciences. Dexamethasone (D4902), Vincristine (V8388), Daunorubicin (30450), Cytarabine (C3350000), and Methotrexate (M9929) were purchased from Millipore Sigma Aldrich. L-asparaginase (RP1792870500) was purchased from BioVendor.

**Cell cycle and cell growth assays**

Cells in logarithmic growth phase were mixed well and separated into two samples after 72-hour drug treatment. Cells were counted by 1:1 mixture of cell suspension with Trypan Blue as a dead cell stain. The remainder of the sample was methanol-fixed, permeabilized, and stained with propidium iodide as a DNA quantity indicator dye and the samples were washed and analyzed by flow cytometry as indicated.

**WST-1 assay**

WST-1 Cell proliferation reagent (5015944001) was purchased from Millipore Sigma Aldrich. Cells were cultured in 96-well plates with escalating doses of drug for 72 hours. 10µL of WST-1 reagent/well was added and the samples were mixed well and incubated at 37 °C for 1 hour. Absorbance was measured on a Perkin Elmer Victor X5 2030 96-well plate reader at 450nm.

#### Supplemental Figures and Table

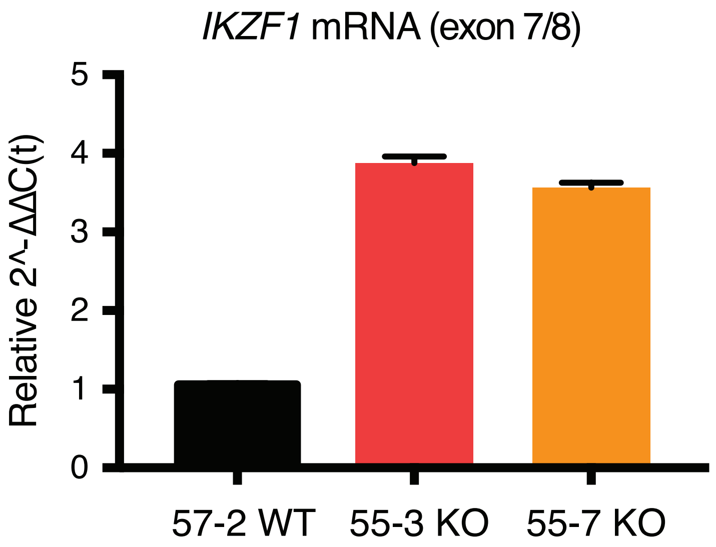


**Supp. Figure S1: *IKZF1* knockout leads to compensatory upregulation of mRNA levels**

qRT-PCR from Nalm-6 clonal cell lines with validated Taqman assay spanning the exon 7-8 junction of *IKZF1*.

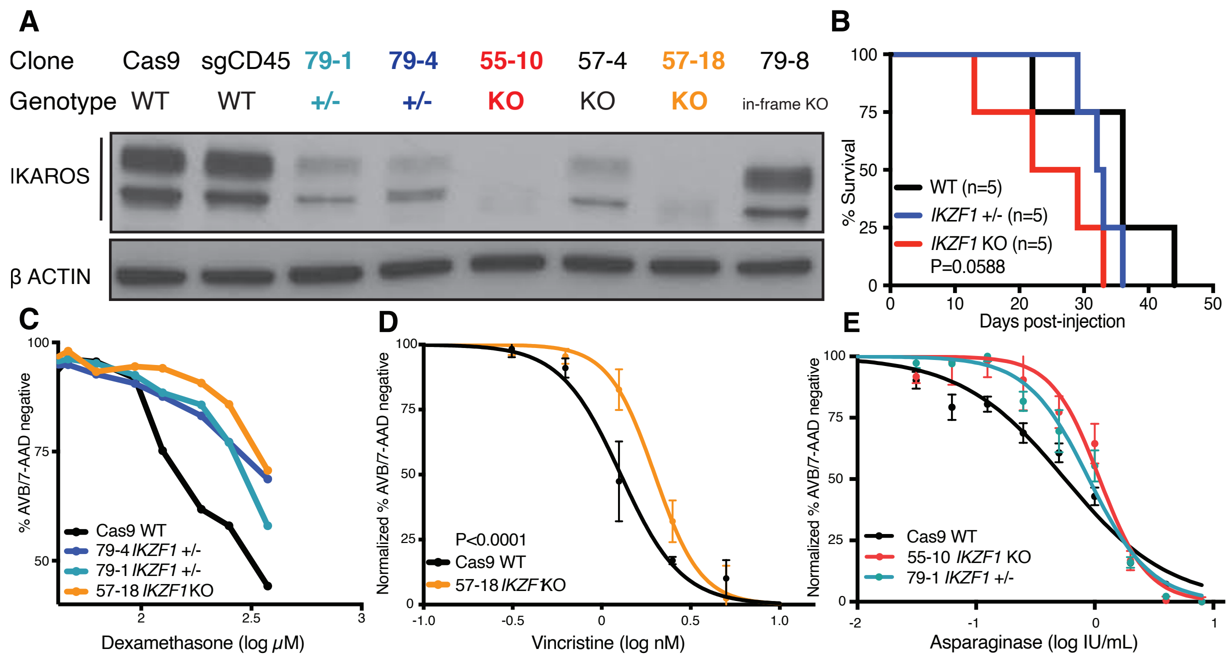


**Supp. Figure S2: *IKZF1* knockout in Tanoue corresponds with Nalm-6 drug resistance and ALL xenograft aggressiveness**

A) IKAROS immunoblotting of single-cell derived Tanoue B-ALL cell line clones with Sanger sequencing-verified *IKZF1* frameshift mutations. Bold coloring of clones indicate those used in further experiments that are similarly color-coded throughout. B) Kaplan-Meier survival curve of NSG mice xenografts transplanted with Tanoue clonal cell lines; n= total number of mice in each group. (C-E) drug treatment of Tanoue clonal cell lines showing percent AnnexinV/7-AAD double negative population as measured by flow cytometry with C) dexamethasone, D) vincristine, and E) asparaginase. Samples were done in triplicate; error bars show SEM. All treatments were 72 hours except for D), 48 hours.

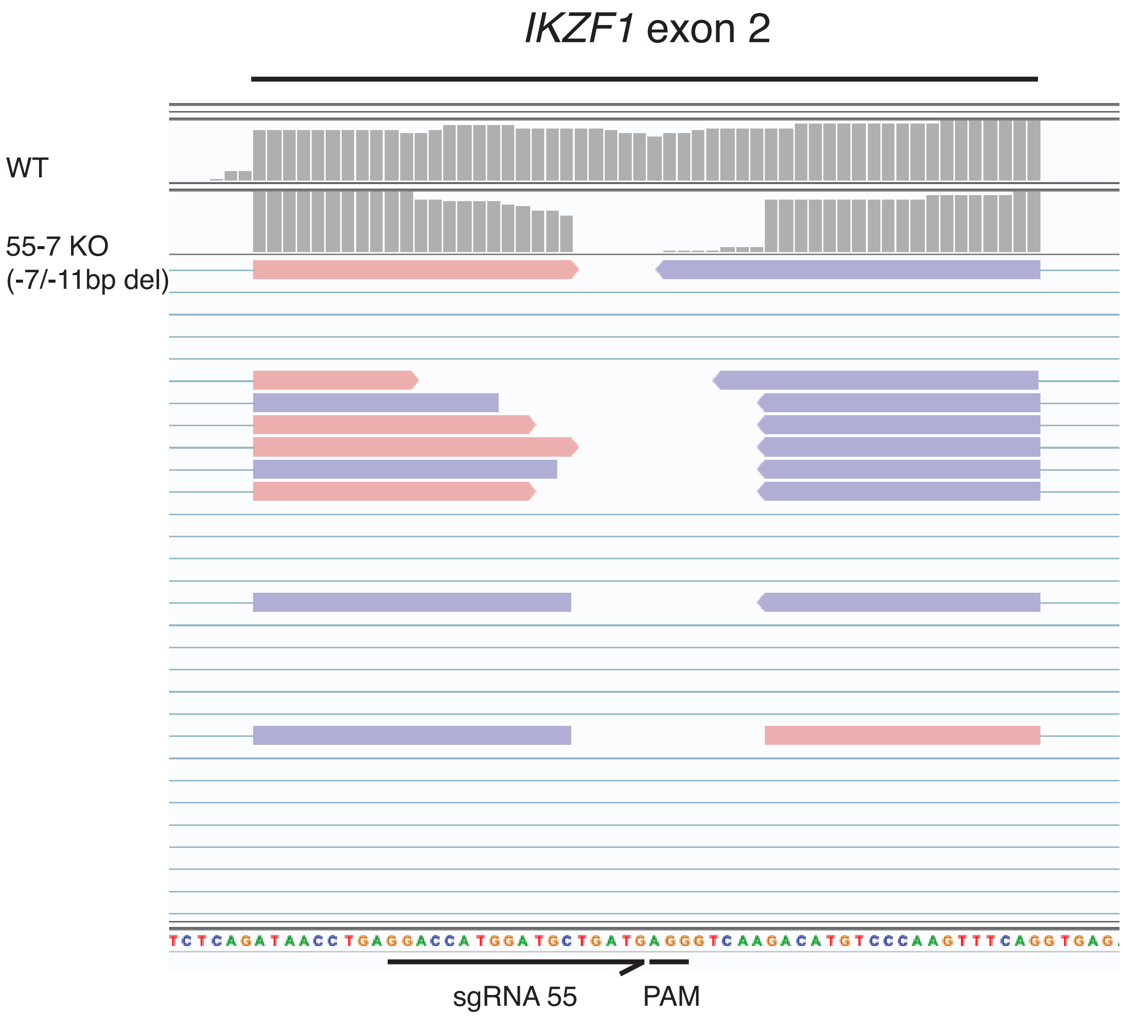


**Supp. Figure S3: RNAseq reads verify of out-of-frame deletion of *IKZF1***

RNAseq reads at targeted exon 2 of *IKZF1* aligned in Integrative Genomics Viewer (IGV) showing predicted biallelic deletions of 7- and 11-base pairs in Nalm-6 clonal cell lines.

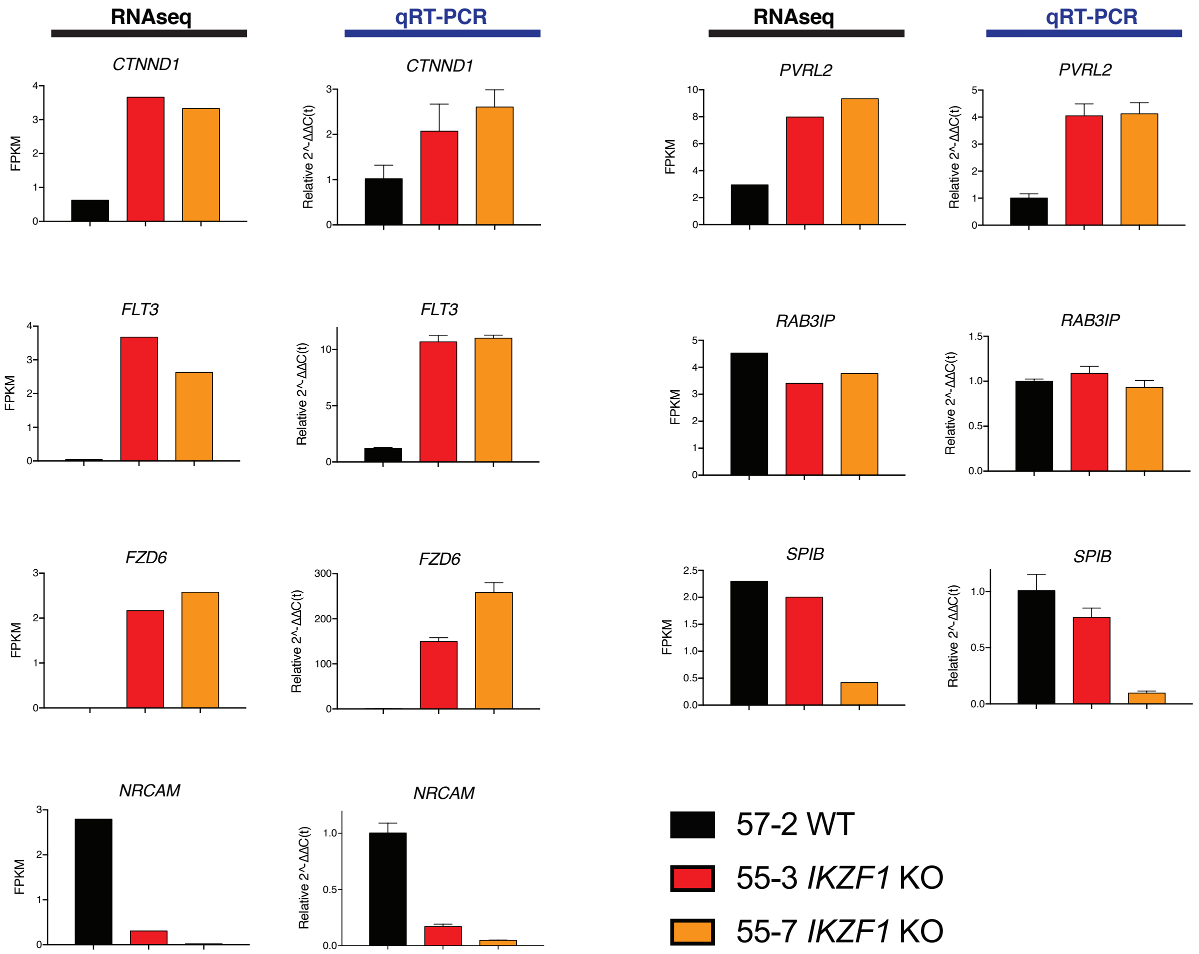


**Supp. Figure S4: qRT-PCR accurately validates RNAseq results with a panel of up- and down-regulated genes after *IKZF1* knockout**

RNAseq in Nalm-6 clonal cell lines of indicated genotype on left (FPKM) and qRT-PCR for the same gene shown on the right (relative expression). Error bars show SEM.

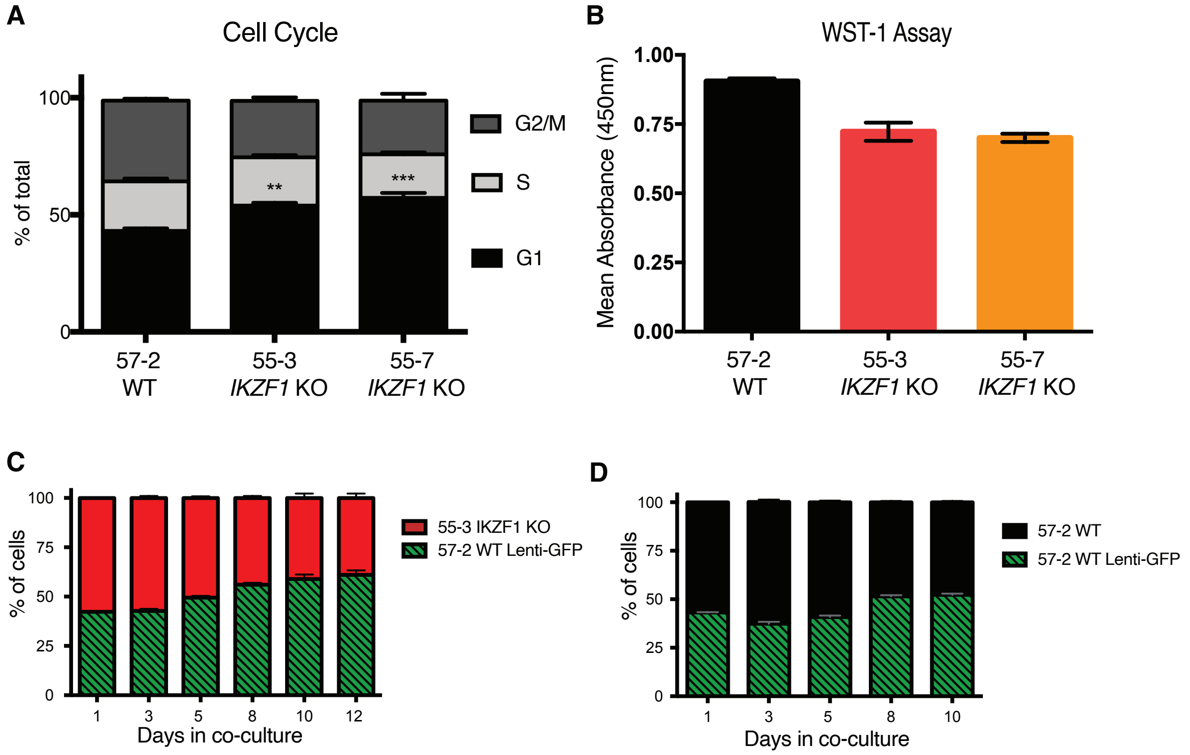


**Supp. Figure S5: *IKZF1* knockout displays lower basal metabolic rate and cell division frequency**

All samples were done in triplicate in Nalm-6 clonal cell lines. Error bars show SEM.

A) Cell cycle analysis with propidium iodide staining for DNA content. B) Baseline WST-1 assay from equal numbers of each cell line. C) WT clone 57-2 transduced with GFP lentivirus was mixed in 1:1 ratio with *IKZF1* KO or D) 57-2 as control and co-cultured for 10-12 days with relative ratio at indicated timepoints measured by flow cytometry for GFP positivity.

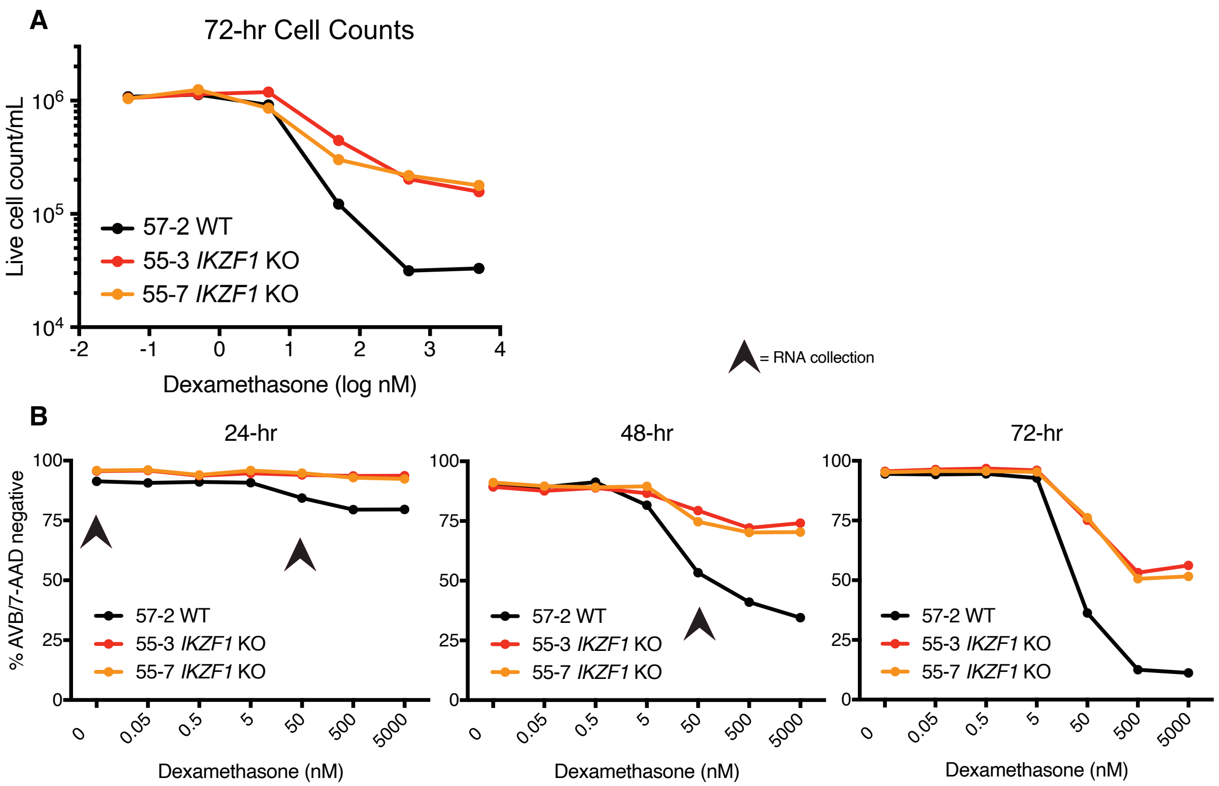


**Supp. Figure S6: Timepoint series of differential glucocorticoid resistance due to *IKZF1* knockout**

Nalm-6 clonal cell lines were subjected to escalating dexamethasone doses. A) automated live-cell counts by trypan blue dead cell exclusion dye after 72-hour treatment. B) percent AnnexinV/7-AAD double negative population as measured by flow cytometry. Arrow symbol indicates all dose-timepoints where RNA was collected from each of three samples for RNAseq.

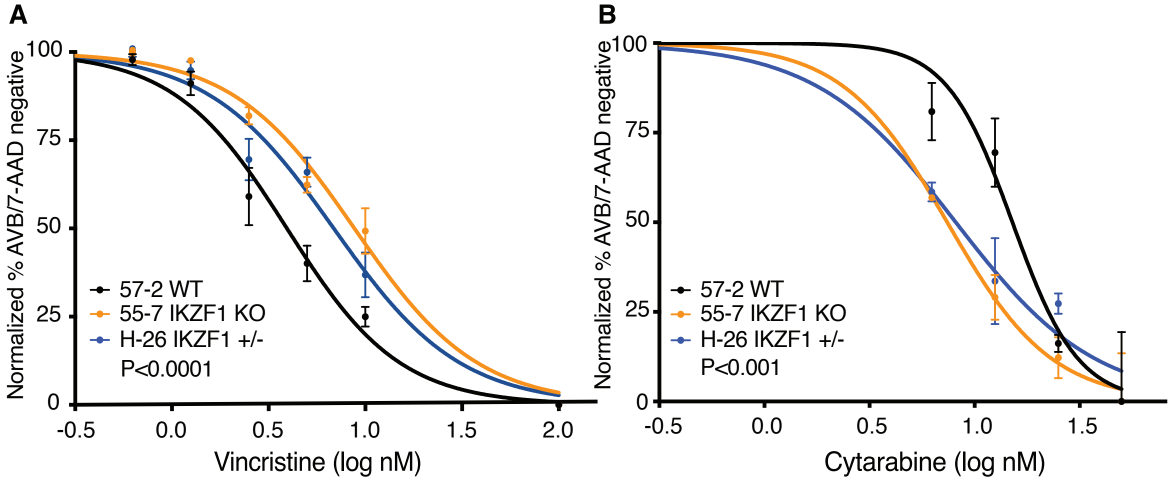


**Supp. Figure S7: Heterozygous deletions of *IKZF1* in Nalm-6 mimic vincristine resistance and cytarabine sensitivity profiles of knockout**

Drug treatment of Nalm-6 clonal cell lines showing percent AnnexinV/7-AAD double negative population as measured by flow cytometry. A) vincristine (48 hours). B) cytarabine (72 hours). Samples were done in triplicate; error bars show SEM.

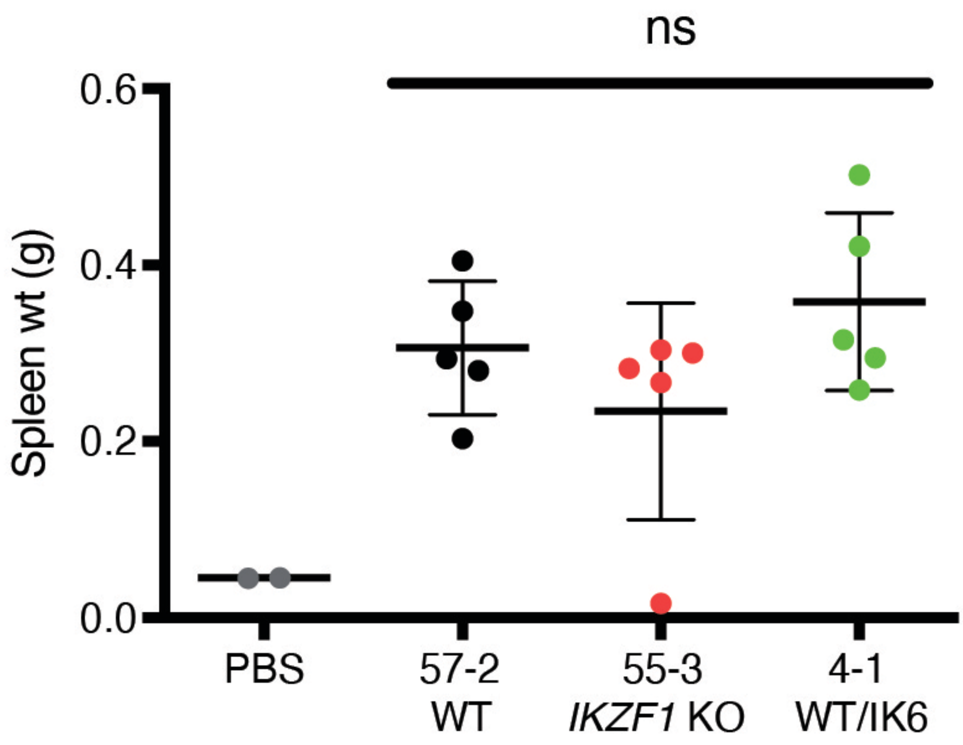


**Supp. Figure S8: Xenograft splenic engraftment is unchanged after *IKZF1* perturbations**

Spleen weight (g) at time of sacrifice of NSG mice xenografts transplanted with indicated Nalm-6 clonal cell lines. ns= not significant. Data points shown correspond to animals from Figure 7A-C. Horizontal line is mean and error bars show SEM.

| Nalm-6 Clone | *IKZF1* Genotype | Dexamethasone $IC_{50}$ (nM) | Cytarabine $IC_{50}$ (nM) | Vincristine $IC_{50}$ (nM) | Daunorubicin $IC_{50}$ (nM) | Asparaginase $IC_{50}$ (IU/mL) | Methotrexate $IC_{50}$ (nM) |
|---|---|---|---|---|---|---|---|
| **57-2** | **WT** | 3.13 | 23.39 | 1.86 | 24.97 | 2.24 | 21.11 |
| **55-3** | **KO** | >5,000 | -- | 2.93 | -- | 5.59 | 22.36 |
| **55-7** | **KO** | >5,000 | 11.62 | 2.73 | 90.25 | 5.53 | 24.42 |
| **4-1** | **WT/IK6** | 10.24 | 8.65 | 3.03 | 38.80 | 2.63 | -- |
| **3-5** | **WT/IK6** | 9.28 | 10.04 | -- | 32.19 | 2.79 | -- |

**Supp. Table S1: Summary of $IC_{50}$ values**

From Nalm-6 clonal cell lines of indicated genotype. $IC_{50}$ values were extrapolated from flow cytometry data using percent AnnexinV/7-AAD double-negative population.

**sgRNA sequences**

| **sgRNA** | **sequence 5'-->3' (sgRNA binding region in CAPS) for IVT template PCR** |
|---|---|
| IKZF1_sgRNA80 | ttaatacgactcactataGGGGTGGTGGAGAGGTCCTCgttttagagctagaaatagc |
| IKZF1_sgRNA79 | ttaatacgactcactataGGGGACCTCTCCACCACCTCgttttagagctagaaatagc |
| IKZF1_sgRNA77 | ttaatacgactcactataGGGGAGAGGTCCTCGGGGATgttttagagctagaaatagc |
| IKZF1_sgRNA75 | ttaatacgactcactataGGTTTGCTGTCCTCCCGAGGgttttagagctagaaatagc |
| IKZF1_sgRNA57 | ttaatacgactcactataGGGACCATGGATGCTGATGAgttttagagctagaaatagc |
| IKZ1F_sgRNA55 | ttaatacgactcactataGGGGACCATGGATGCTGATGgttttagagctagaaatagc |

**Primers**

| **PCR/Sanger sequencing Primers** | **sequence 5'-->3'** |
|---|---|
| hIKZF1_EX2_F | ggatcaaggtctgtgccagtctga |
| hIKZF1_EX2_R | cacttcctgggtctcctggatgag |
| hIKZF1_EX3_F | aagagtggcagtgtctgggattat |
| hIKZF1_EX3_R | caacttggacattttgcggggtg |
| F 5' long arm int2 EcoRI | agtcgggaattccagaaagcaatgtggac |
| R 5' long arm amplify | gagcaccgcttccacatgagc |
| F GFP 3' arm amplify | gccaggaccggtacgagttct |
| R 3' arm amplify EcoRI | gctcaagaattcagagggataacataaaa |

**qRT-PCR Taqman probes**

| **Target gene** | **Taqman Probe Catalog Number: ThermoFisher/Applied Biosystems** |
|---|---|
| *CTNND1* | Hs00931670 |
| *FLT3* | Hs00174690 |
| *FZD6* | Hs00171574 |
| *GAPDH* | Hs02786624 |
| *IKZF1* exon 7-8 | Hs00958479 |
| *NRCAM* | Hs01031598 |
| *NECTIN2 (PVRL2)* | Hs01071562 |
| *RAB3IP* | Hs00223789 |
| *SPIB* | Hs00162510 |

#### HDR Template Sequences (used with sgRNA 79)

Note: The two 5 prime templates vary only by codon optimization in Exon 3 between the sgRNA cleavage site and Exon8 junction in the 2nd template labeled 'cod'.
The templates were PCR amplified singly, AgeI digested, ligated and the entire fragment was then apmlified again.

| **HDR Template for Ik6 Knock-in** | **Description** | **sequence 5'-->3'** |
|---|---|---|
| IK-6_GFP_ mutSG_5pri | EcorI 5 prime arm int2 ex3 ex8 mutated sg site only | AGTCGGGAATTCCAGAAAGCAATGTGGACTCAAAAATAGGAAAAGAATTTAAATAAATCCTAGTGTACCCGTTATACATGAAA<br>TTATGGAAATATGACCCTGAGCATGGAAATATGTACATGAGAATGTCTAAAAGCTAGTTCATTTTGAAAAACAAAATAATGTC<br>ACCTCATATTATTTATAGTATATAAAGATGATTTTAAGAGTGGCAGTGTCTGGGATTATAGGTGATTGTATTTCTTCCCTTTT<br>GCACATCTATGTTCTCTCATTTGTATTGTGTGGGGAGAAGTGACTTTTTTTATAAAAAGAAAAAGGTATATGCATCCCAGCAG<br>AGAAGCACTGGCTCCACCCAGTACCTGCCTCCTCATGCCACCCTCTCAAGCCAAAAGCCGGGGGAAGCCCAGGCACCTTGACC<br>ATGACCGCCCGAGACTCACACTTCTTCTTTCTCATCAGGGAAGGAAAGCCCCCCTGTAAGCGATACTCCAGATGAGGGCGATG<br>AGCCCATGCCGATCCCCGAGGACCTCTCCACGACGTCGGGAGGACAGCAAAGCTCCAAGAGTGACAGAGTCGTGGGGGACAAG<br>GGCCTGTCCGACACGCCCTACGACAGCAGCGCCAGCTACGAGAAGGAGAACGAAATGATGAAGTCCCACGTGATGGACCAAGC<br>CATCAACAACGCCATCAACTACCTGGGGGCCGAGTCCCTGCGCCCGCTGGTGCAGACGCCCCCGGGCGGTTCCGAGGTGGTCC<br>CGGTCATCAGCCCGATGTACCAGCTGCACAAGCCGCTCGCGGAGGGCACCCCGCGCTCCAACCACTCGGCCCAGGACAGCGCC<br>GTGGAGAACCTGCTGCTGCTCTCCAAGGCCAAGTTGGTGCCCTCGGAGCGCGAGGCGTCCCCGAGCAACAGCTGCCAAGACTC<br>CACGGACACCGAGAGCAACAACGAGGAGCAGCGCAGCGGTCTCATCTACCTGACCAACCACATCGCCCCGCACGCGCGCAACG<br>GGCTGTCGCTCAAGGAGGAGCACCGCGCCTACGACCTGCTGCGCGCCGCCTCCGAGAACTCGCAGGACGCGCTCCGCGTGGTC<br>AGCACCAGCGGGGAGCAGATGAAGGTGTACAAGTGCGAACACTGCCGGGTGCTCTTCCTGGATCACGTCATGTACACCATCCA<br>CATGGGCTGCCACGGCTTCCGTGATCCTTTTGAGTGCAACATGTGCGGCTACCACAGCCAGGACCGGTACGAGTTCTCGTCGC<br>ACATAACGCGAGGGGAGCACCGCTTCCACATGAGC |
| IK-6_GFP_ mutSG-ex3cod_5pri | EcoRI 5 prime arm int2 ex3 ex8 mutated sg site cod opt ex3 | AGTCGGGAATTCCAGAAAGCAATGTGGACTCAAAAATAGGAAAAGAATTTAAATAAATCCTAGTGTACCCGTTATACATGAAA<br>TTATGGAAATATGACCCTGAGCATGGAAATATGTACATGAGAATGTCTAAAAGCTAGTTCATTTTGAAAAACAAAATAATGTC<br>ACCTCATATTATTTATAGTATATAAAGATGATTTTAAGAGTGGCAGTGTCTGGGATTATAGGTGATTGTATTTCTTCCCTTTT<br>GCACATCTATGTTCTCTCATTTGTATTGTGTGGGGAGAAGTGACTTTTTTTATAAAAAGAAAAAGGTATATGCATCCCAGCAG<br>AGAAGCACTGGCTCCACCCAGTACCTGCCTCCTCATGCCACCCTCTCAAGCCAAAAGCCGGGGGAAGCCCAGGCACCTTGACC<br>ATGACCGCCCGAGACTCACACTTCTTCTTTCTCATCAGGGAAGGAAAGCCCCCCTGTAAGCGATACTCCAGATGAGGGCGATG<br>AGCCCATGCCGATCCCCGAGGACCTCTCCACGACGTCGGGAGGGCAACAGAGCAGTAAGTCTGATAGGGTGGTGGGGGACAAG<br>GGCCTGTCCGACACGCCCTACGACAGCAGCGCCAGCTACGAGAAGGAGAACGAAATGATGAAGTCCCACGTGATGGACCAAGC<br>CATCAACAACGCCATCAACTACCTGGGGGCCGAGTCCCTGCGCCCGCTGGTGCAGACGCCCCCGGGCGGTTCCGAGGTGGTCC<br>CGGTCATCAGCCCGATGTACCAGCTGCACAAGCCGCTCGCGGAGGGCACCCCGCGCTCCAACCACTCGGCCCAGGACAGCGCC<br>GTGGAGAACCTGCTGCTGCTCTCCAAGGCCAAGTTGGTGCCCTCGGAGCGCGAGGCGTCCCCGAGCAACAGCTGCCAAGACTC<br>CACGGACACCGAGAGCAACAACGAGGAGCAGCGCAGCGGTCTCATCTACCTGACCAACCACATCGCCCCGCACGCGCGCAACG<br>GGCTGTCGCTCAAGGAGGAGCACCGCGCCTACGACCTGCTGCGCGCCGCCTCCGAGAACTCGCAGGACGCGCTCCGCGTGGTC<br>AGCACCAGCGGGGAGCAGATGAAGGTGTACAAGTGCGAACACTGCCGGGTGCTCTTCCTGGATCACGTCATGTACACCATCCA<br>CATGGGCTGCCACGGCTTCCGTGATCCTTTTGAGTGCAACATGTGCGGCTACCACAGCCAGGACCGGTACGAGTTCTCGTCGC<br>ACATAACGCGAGGGGAGCACCGCTTCCACATGAGC |
| IK-6_GFP_ sv40_3pri | 3 prime arm GFP + sv40 int 3 ecorI | GCCAGGACCGGTACGAGTTCTCGTCGCACATAACGCGAGGGGAGCACCGCTTCCACATGAGCATGGTGAGCAAGGGCGAGGAG<br>CTGTTCACCGGGGTGGTGCCCATCCTGGTCGAGCTGGACGGCGACGTAAACGGCCACAAGTTCAGCGTGTCCGGCGAGGGCGA<br>GGGCGATGCCACCTACGGCAAGCTGACCCTGAAGTTCATCTGCACCACCGGCAAGCTGCCCGTGCCCTGGCCCACCCTCGTGA<br>CCACCCTGACCTACGGCGTGCAGTGCTTCAGCCGCTACCCCGACCACATGAAGCAGCACGACTTCTTCAAGTCCGCCATGCCC<br>GAAGGCTACGTCCAGGAGCGCACCATCTTCTTCAAGGACGACGGCAACTACAAGACCCGCGCCGAGGTGAAGTTCGAGGGCGA<br>CACCCTGGTGAACCGCATCGAGCTGAAGGGCATCGACTTCAAGGAGGACGGCAACATCCTGGGGCACAAGCTGGAGTACAACT<br>ACAACAGCCACAACGTCTATATCATGGCCGACAAGCAGAAGAACGGCATCAAGGTGAACTTCAAGATCCGCCACAACATCGAG<br>GACGGCAGCGTGCAGCTCGCCGACCACTACCAGCAGAACACCCCCATCGGCGACGGCCCCGTGCTGCTGCCCGACAACCACTA<br>CCTGAGCACCCAGTCCGCCCTGAGCAAAGACCCCAACGAGAAGCGCGATCACATGGTCCTGCTGGAGTTCGTGACCGCCGCCG<br>GGATCACTCTCGGCATGGACGAGCTGTACAAGTAAATCTAGATAACTGATCATAATCAGCCATACCACATTTGTAGAGGTTTT<br>ACTTGCTTTAAAAAACCTCCCACACCTCCCCCTGAACCTGAAACATAAAATGAATGCAATTGTTGTTGTTAACTTGTTTATTG<br>CAGCTTATAATGGTTACAAATAAAGCAATAGCATCACAAATTTCACAAATAAAGCATTTTTTTCACTGCATTCTAGTTGTGGT<br>TTGTCCAAACTCATCAATGTATCTTAGTAAGTGGGTCACCAGCGGCCTCTGTGCCTGTGAAACCTTTATCTCTTTGTATTTTT<br>CCAAGACAGTGATGAAGGGATGCAAGTCATTTTATCCATTGTGTTCCCTCAACTGGCATATTAAAGAAATATGGCACAAAGAT<br>CAGCAGGATGGGGGTCCTCTGGTGTGTGGGAGGATGGACACTCACAGGCCAGCATGGCCGTGAGAGCCACACACCCCGCAAAA<br>TGTCCAAGTTGAGGAGCAATCCTGCCCAGGGACGCGTCTCTGTCACTGTCCTCTGTCCTCACTGCACTTGCAGGAATATCAAA<br>TGTTATGGATTGTAGATCGTGAAAATTACACACTTACGTGTTTTGGCAACAGTGCTTTTCAGTGTTTGGGGTTAGAACAAGCC<br>ACATCTGGCCATTTTATGTTATCCCTCTGAATTCTTGAGC |
